## Supplementary Figures for "Endogenous fluctuations of OCT4 and SOX2 bias pluripotent cell fate decisions"

**Supplementary Figure 1: Validation of knock-in cell lines**

**a** General knock-in strategy used in this study. Red hexagons: STOP codons; KIC: knock-in construct. **b** Location of primers (arrows) and PCR results to verify knocked-in reporters in the SNSF and SBROS cell lines. Red hexagon: STOP codons; Yellow triangles: LoxP sites. **c** Western blot confirming the tagging of endogenous SOX2 and OCT4 in the SNSF and SBROS cell lines. **d** Example of field of view used for the correlation analysis of total OCT4 ($\alpha$OCT4) and heterozygous OCT4-HALO. Scale bar: 50µm. **e** QPCR analysis of pluripotency marker expression for cell lines used in this study, normalized to the levels of CGR8 wt mouse ES cells (n=3). **f** Representative images of Alkaline Phosphatase staining in the different cell lines. **g** Ratio of the number of colonies with versus without dox, for wt, Ypet-OCT4 or OCT4-Halo-expressing Zhbtc4 cells (n=3). **h** Co-immunoprecipitation (Co-IP) of OCT4 and OCT4-HALO with SOX2-SNAP (SBROS) and OCT4 with wt SOX2 (SBR). Arrow: Samples and molecular weight marker (MWM) were spliced together. **i** Heatmaps for SOX2 (GSE89599) and YPet-SOX2 ChIP-seq signals in SOX2-bound regions and OCT4, OCT4-HALO and YPet-OCT4 ChIP-seq signals in OCT4-bound regions. **j** Ratio of dome-shaped/flat colonies for wt or YPet-SOX2-expressing 2TS22C cells in the absence or presence of dox (n=3). All p-values were determined using a two-sided t-test; Error bars: SE.

**Supplementary Figure 2: Cross regulation of SOX2 and OCT4**

**a-c** YPET-OCT4 overexpression time-course. **a** fluorescence microscopy images after immunofluorescence staining of SOX2 and OCT4. OE: overexpressed YPET-OCT4. **b,c** quantifications of total SOX2 and OCT4 levels in single cells upon YPET-OCT4 overexpression, normalized to the average at 0h. Whiskers: minimum and maximum values; Box: lower and upper quartiles; Solid line: median; Outliers: solid points. **d-f** YPET-SOX2 overexpression time-course. **d** fluorescence microscopy images after immunofluorescence staining of SOX2 and OCT4. OE: overexpressed YPET-SOX2. **e,f** quantifications of total SOX2 and OCT4 levels in single cells upon YPET-SOX2 overexpression, normalized to the average at 0h. **f** induction efficiency of YPET-SOX2 upon dox addition. Whiskers: minimum and maximum values; Box: lower and upper quartiles; Solid line: median; Outliers: solid points. **g** Total OCT4 or SOX2 levels as a function of YPET-OCT4 or YPET-SOX2 overexpression, respectively, 4 hours (blue) and 7 hours (red) after addition of dox. **h** Scheme of the knock-in alleles in the SNSF cell line (Red hexagons: STOP codons; Yellow triangles: LoxP sites) and luminescence microscopy images of differentiating SNSF cells showing the SOX2-NLUC and SOX1-P2A-FLUC signal (Scale bar: 50µm). **i** Distributions of OCT4 (n=2682), SOX2 (n=1236) and NANOG (n=2416) levels in wt E14 and SNSF cell lines as determined by quantitative immunofluorescence. Dashed lines: means protein levels; Dotted lines median protein levels. **j** Cell cycle duration of E14 wt (n=6) and SNSF (n=6) cells. Error bars: SE. p value: two-sided t-test. **k** Strategy to sort G1 cells for medium endogenous levels of SOX2 and high or low OCT4 levels (top), and conversely (bottom). **l** Raw data of SOX2 and OCT4 levels 8h after sorting as indicated in panel k. Scale-bars: 50$\mu$m.

**Supplementary Figure 3: Fluctuations of SOX2 and OCT4 over the cell cycle**

**a-b** Average SOX2 protein level (**a**) and concentration (**b**) (n=164, shaded area: SD) in SNSF cells. **c** Changes of single cell ranks of SOX2 levels over time using data from one cell cycle (n=100). Red: initially high expressing cells; Blue: initially low expressing cells. **d** Rank-based autocorrelation function of the SOX2 ranks. Error bars: SE estimated by bootstrapping. **e-f** Average OCT4 protein level (**e**) and concentration (**f**) (n=48) in SBROS cells; shaded area: SD.

**Supplementary Figure 4: Impact of SOX2 and OCT4 levels on differentiation outcomes**

**a,b** Sorting strategies to evaluate the impact of SOX2 (**a**) or OCT4 (**b**) levels on differentiation outcomes and example of flow cytometry data after four days of differentiation. **c** Example of flow cytometry data after four days of differentiation with or without OCT4 overexpression for 12 hours before differentiation. **d** Example of flow cytometry data after four days of differentiation with or without dox throughout differentiation. **e** Differentiation outcome after four days with or without 100ng/ml dox throughout differentiation. % of SOX1+ and BRA+ cells are normalized on the average values of each experiment (n=6). p-values were determined using a two-sided t-test. **f** Sorting strategy for SHOH, SHOL, SLOH and SLOL populations in G1-phase. **g** Example of flow cytometry data of SHOH, SHOL, SLOH and SLOL sorted in G1 phase (see Fig.3i). **h** Average fraction of BRA+ and SOX1+ cells after four days of differentiation, when sorted for G1 or S- phase, respectively (n=5). p-values were determined using a two-sided t-test. **i** Example of flow cytometry data of SHOH, SHOL, SLOH and SLOL sorted in S-phase.

**Supplementary Figure 5: ATAC-seq analysis, GO Enrichment Analysis and phenotype of Eomes enhancer mutants**

**a** Fraction of reads in ATAC-seq peaks for the different cell populations. **b** Violin plot of OCT4-high vs OCT4-low and SOX2-high vs SOX2-low in different groups of loci. **c** Metaplots of chromatin accessibility for the different cell populations in different groups of loci. **d-h** Top 12 GO terms of loci in the Upregulated SOX2 (**d**), Upregulated OCT4&SOX2 (**e**), Downregulated OCT4 (**f**), Downregulated SOX2 (**g**), and Downregulated OCT4&SOX2 (**h**) groups. **i** Top 12 GO terms for loci in the Upregulated OCT4 group that are bound by OCT4 and with FDR < 5%. **j** Examples of normalized tag counts in enhancers close to Rbpj (top) and Eomes (bottom) in the Upregulated OCT4 group for different replicates. **k** Heatmaps for OCT4 (GSE87820 and GSE56138) and SOX2 (GSE87820) in different groups of loci. **l** Changes in accessibility of regions up or down- regulated in OCT4-high cells upon OCT4 knock-down (GSE87822).

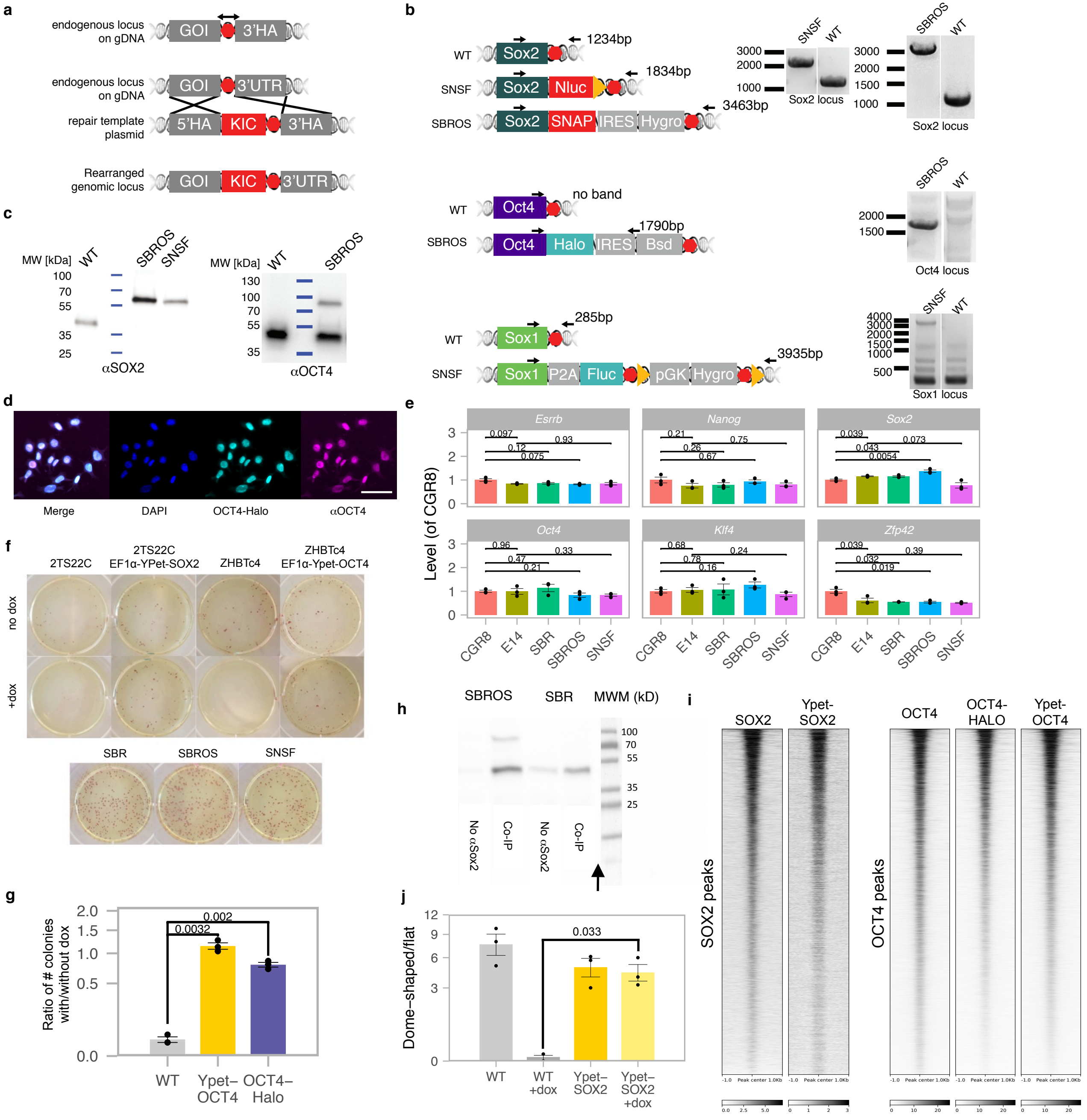


Supplementary Figure 1

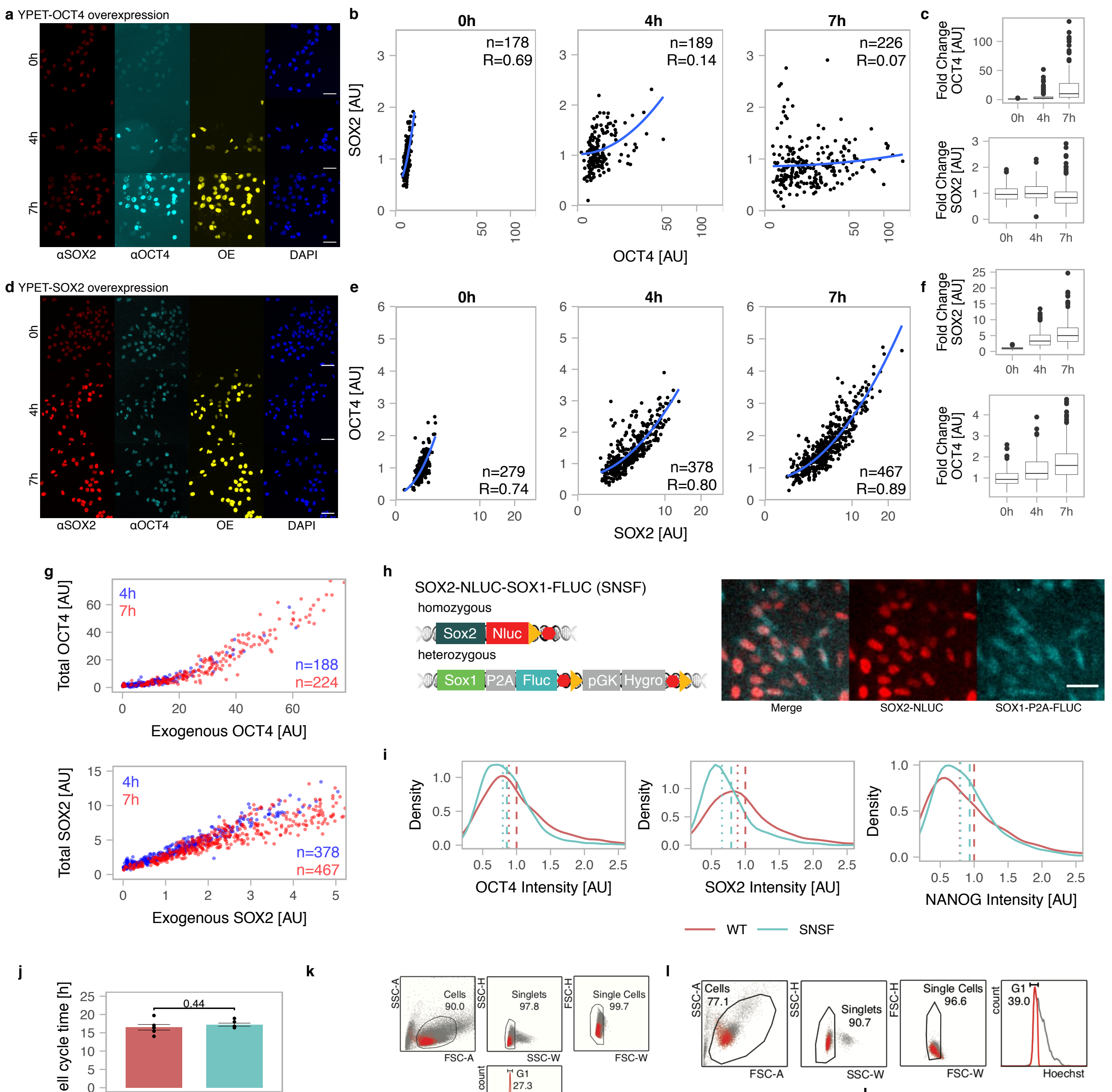


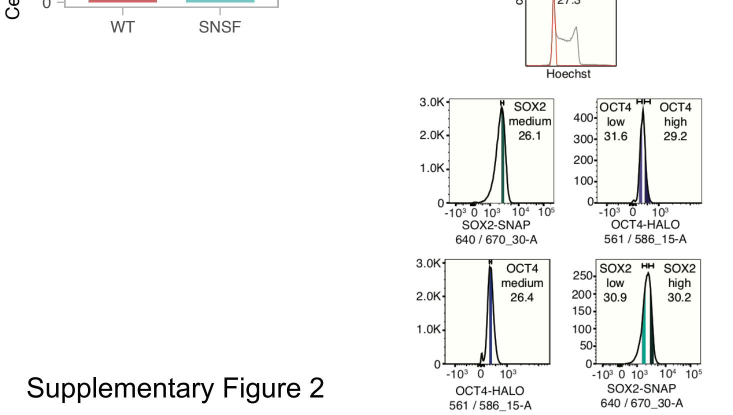


| | N2B27+2iLIF SOX2 [AU] | N2B27+2iLIF OCT4 [AU] | N2B27 SOX2 [AU] | N2B27 OCT4 [AU] |
|---|---|---|---|---|
| SOX2 high | 18570 | 10677 | 19255 | 10649 |
| | 37134 | 22004 | 39463 | 21126 |
| | 38511 | 17281 | 34713 | 15074 |
| | 8376 | 11430 | 7714 | 12914 |
| SOX2 low | 12437 | 6769 | 12433 | 6782 |
| | 32140 | 20656 | 27365 | 18453 |
| | 29544 | 15502 | 25516 | 15642 |
| | 5589 | 12097 | 5275 | 12232 |
| OCT4 high | 17800 | 11809 | 18107 | 12575 |
| | 37848 | 21981 | 31363 | 19771 |
| | 35557 | 20283 | 32022 | 18668 |
| | 6160 | 12462 | 5698 | 13029 |
| OCT4 low | 14752 | 6425 | 14777 | 7247 |
| | 34152 | 19830 | 37070 | 18334 |
| | 36378 | 21418 | 31604 | 15006 |
| | 7514 | 8435 | 7100 | 9614 |

Supplementary Figure 2

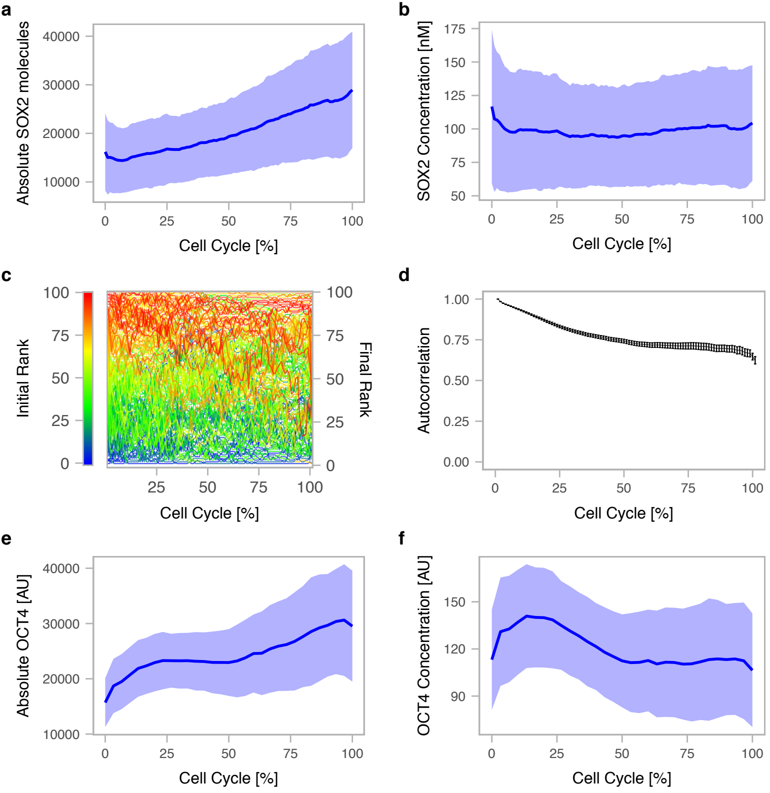


Supplementary Figure 3

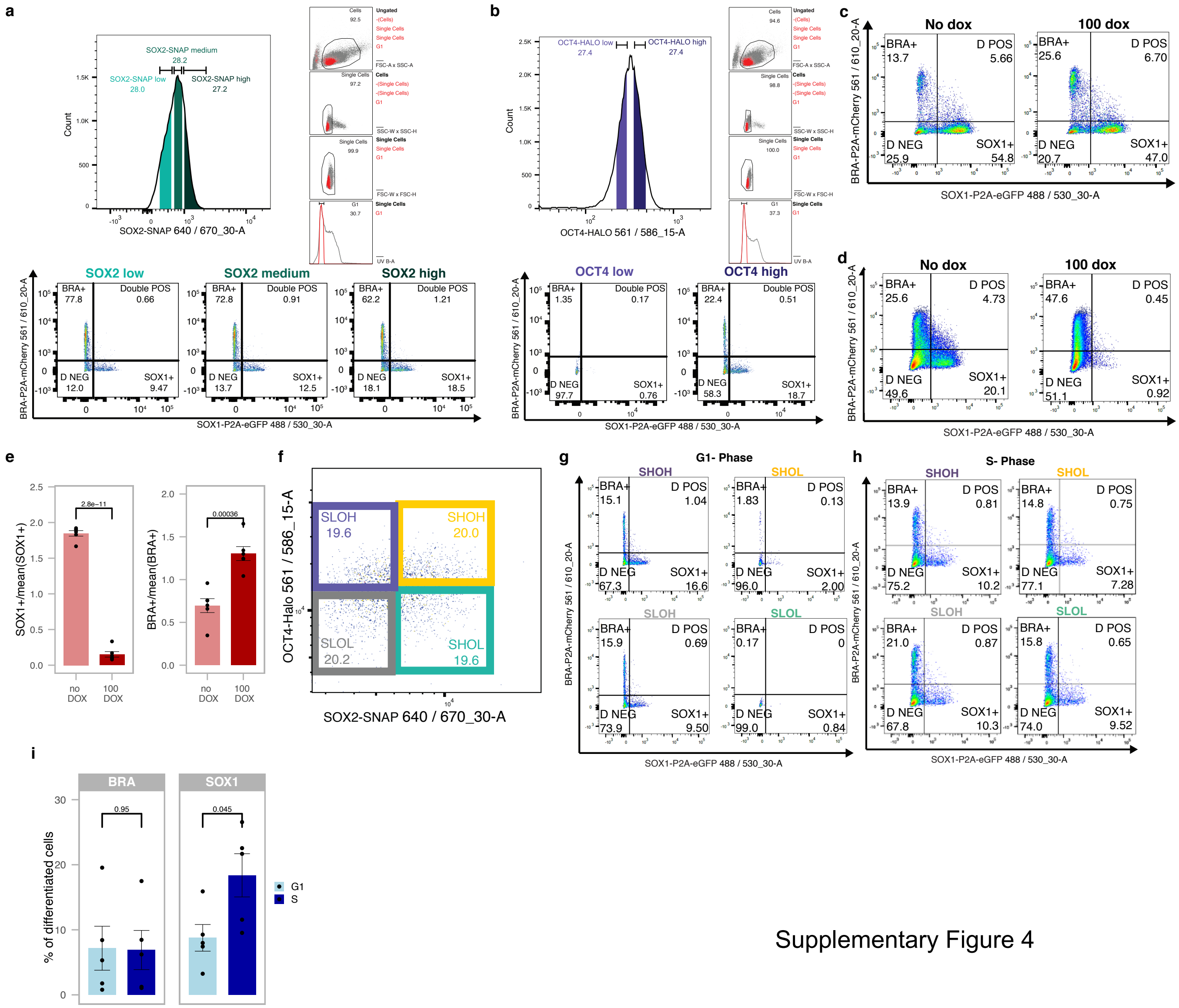


Supplementary Figure 4

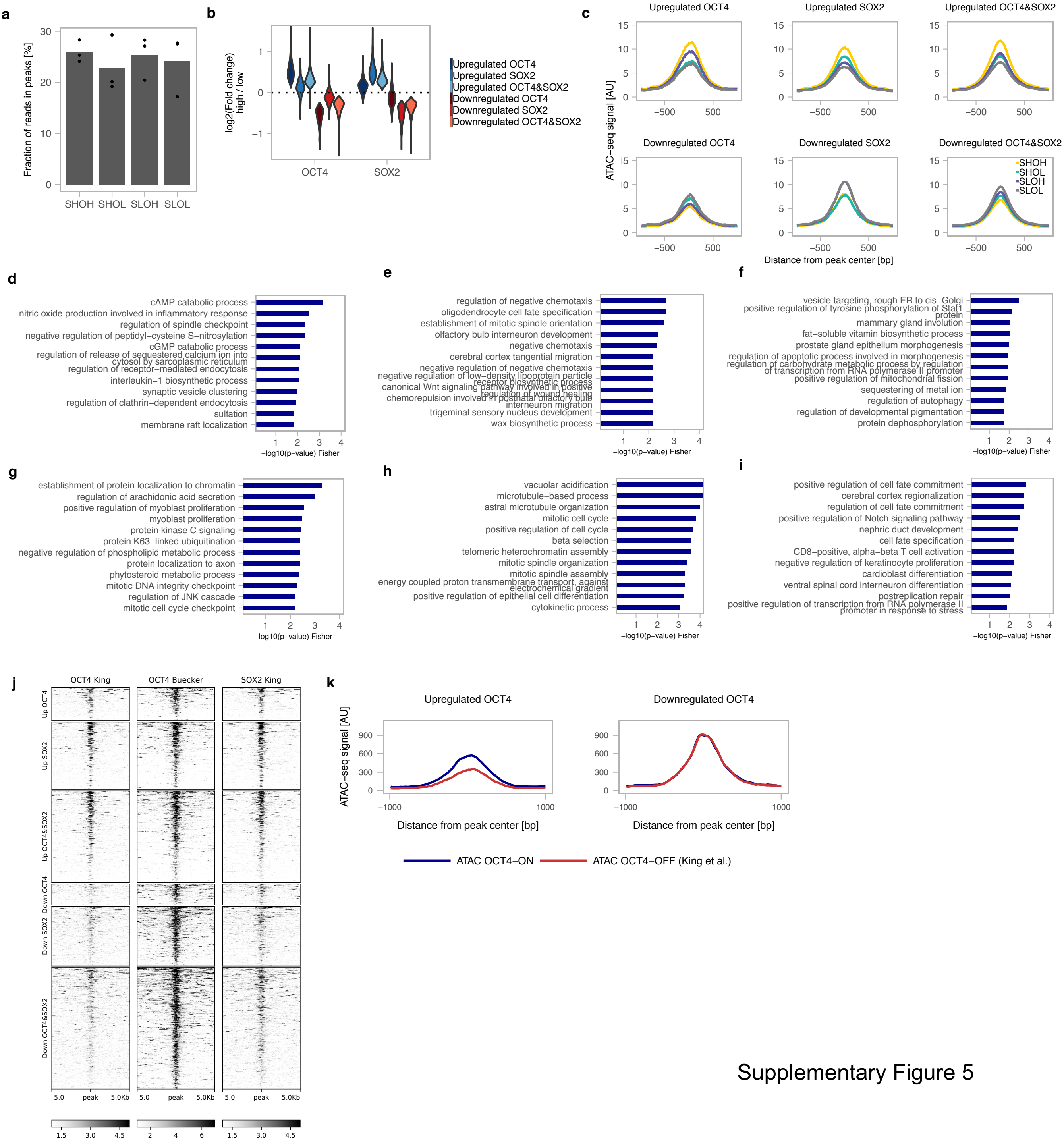


Supplementary Figure 5
